# Investigating higher order interactions in single cell data with scHOT

**DOI:** 10.1101/841593

**Authors:** Shila Ghazanfar, Yingxin Lin, Xianbin Su, David M. Lin, Ellis Patrick, Ze Guang Han, John C. Marioni, Jean Yee Hwa Yang

**Affiliations:** Cancer Research UK Cambridge Institute, University of Cambridge, Cambridge, United Kingdom; School of Mathematics and Statistics, The University of Sydney, Sydney, New South Wales, 2006, Australia; Key Laboratory of Systems Biomedicine (Ministry of Education) and Collaborative Innovation Center of Systems Biomedicine, Shanghai Center for Systems Biomedicine, Shanghai Jiao Tong University, Shanghai 200240, China; Department of Biomedical Sciences, Cornell University, Ithaca, United States; Westmead Institute for Medical Research, Sydney, New South Wales, 2145, Australia; EMBL European Bioinformatics Institute, Wellcome Trust Genome Campus, Cambridge, United Kingdom; Charles Perkins Centre, The University of Sydney, Sydney, New South Wales, 2006, Australia

## Abstract

Single-cell RNA-sequencing has transformed our ability to examine cell fate choice. For example, in the context of development and differentiation, computational ordering of cells along ‘pseudotime’ enables the expression profiles of individual genes, including key transcription factors, to be examined at fine scale temporal resolution. However, while cell fate decisions are typically marked by profound changes in expression, many such changes are observed in genes downstream of the initial cell fate decision. By contrast, the genes directly involved in the cell fate decision process are likely to interact in subtle ways, potentially resulting in observed changes in patterns of correlation and variation rather than mean expression prior to cell fate commitment. Herein, we describe a novel approach, scHOT – single cell Higher Order Testing - which provides a flexible and statistically robust framework for identifying changes in higher order interactions among genes. scHOT is general and modular in nature, can be run in multiple data contexts such as along a continuous trajectory, between discrete groups, and over spatial orientations; as well as accommodate any higher order measurement such as variability or correlation. We demonstrate the utility of scHOT by studying embryonic development of the liver, where we find coordinated changes in higher order interactions of programs related to differentiation and liver function. We also demonstrate its ability to find subtle changes in gene-gene correlation patterns across space using spatially-resolved expression data from the mouse olfactory bulb. scHOT meaningfully adds to first order effect testing, such as differential expression, and provides a framework for interrogating higher order interactions from single cell data.

## INTRODUCTION

Understanding the mechanisms that underpin cell fate choices is a key challenge in developmental biology. It requires disentangling the complex interplay between cell autonomous factors, such as gene expression, and non-autonomous factors such as the signaling environment. In the former context, recent technological advances have enabled the rapid and high-throughput measurement of mRNA expression levels in individual cells. Such single-cell RNA-sequencing (scRNA-seq) datasets have facilitated the generation of atlases of cell types during development in human, mouse, zebrafish and the frog^1–5^. Using such data, cells can be computationally ordered along ‘pseudotime’ and changes in the expression profiles of individual genes can be subsequently determined. However, while cell fate decisions are typically associated with profound changes in expression, many such changes are downstream of the initial cell fate decision. Instead, subtle changes in patterns of variation and coexpression of genes across developmental time, sometimes not associated with substantial changes in mean expression, have been argued to play a more critical role in symmetry breaking^6,7^. Consistent with this, higher order interactions (i.e., looking beyond changes in mean expression) have proved highly informative for understanding genomics data, for example in supervised machine learning settings^8^ and for estimation of unknown spatial patterning^9^. Additionally, with recent developments in high-throughput and high-resolution spatially resolved gene expression mapping (e.g., Spatial Transcriptomics^10^; seqFISH^11^; MERFISH^12^) it is now possible to explore the relationship between higher-order interactions and spatial location. For example, in the context of embryogenesis, do small numbers of spatially-localized cells display aberrantly higher variability in expression profiles prior to committing to a downstream fate?

From a computational perspective, methods for studying higher-order interactions are currently lacking. Although numerous methods have been developed for ordering cells along pseudotime, a computationally derived prediction of cell-type differentiation trajectories^13–17^, methods for identifying individual genes that significantly change their expression levels across the pseudotemporal trajectory^18–20^ typically focus on changes in mean expression of single genes and do not characterize subtle changes in patterns of covariation between subsets of genes across this trajectory. In those cases where higher-order interactions have been studied, a typical analysis aiming to compare correlation patterns along pseudotime first defines strict nonoverlapping sets of cells before estimation of a covariance network, either through direct thresholding on the correlation matrix or using other methods^21,22^. However, estimation of such networks is noisy^23^, and ignores potentially subtle but consistent changes across a continuum, as well as requiring an often ad hoc dichotomization or classification of cells into discrete groups. As we have previously discussed^24^, treating the sample ranking as a covariate and testing for an interaction effect in a linear model is restricted to identifying linear and thus monotonic interactions, which may not be present, especially in highly dynamic or complex trajectories going through multiple changes in the differentiation process. In the context of spatially resolved gene expression data, fewer methods exist, with the focus being on testing the existence of pre-defined patterns^25^ (e.g., a signaling gradient); however, these require *a priori* knowledge about the spatial structures of interest.

Here we introduce single cell Higher Order Testing (scHOT), a framework for examining changes in higher order structure, such as correlation among genes across differentiation pseudotime, among discrete groups, and across spatial landscapes. scHOT builds on our previous work, DCARS (Differential Correlation Across Ranked Samples), which used bulk RNA-sequencing data to test for changes in gene-gene correlation across ranked individual samples^24^. Our approach requires one of the following types of cell-specific information (Figure 1): A) a ranking of cells, which will typically be across pseudotime, or B) spatial coordinates in either two or three dimensions. In the case where spatial coordinates are inferred^26^, scHOT is also applicable using either the cell ranking along a gradient or in the inferred 2/3D space. Given this cell-specific information, as well as a scheme for determining local sample-specific weights, we calculate local higher order interaction vectors among single genes or pairs of genes, uncovering local changes in variability or covariation respectively (Figure 1). Sample-wise permutation testing is then used to assess statistical significance, while retaining the global variability or correlation structure of the original data. This framing of the significant genes and gene-pairs in terms of the set of local higher order interaction estimates allows patterns of changes across the trajectory, groups, or space to be characterized in terms of the higher order interaction, rather than simply by changes in the mean. Moreover, scHOT identifies groups of genes for which similar higher order patterns arise. For a more detailed discussion of how scHOT can be applied in practice, see the Supplementary Note. We demonstrate the utility of scHOT by studying the inferred trajectories of mouse liver hepatoblasts into structural cholangiocytes or functional liver hepatocytes, and illustrate its generalizability in a spatial setting by performing higher order testing on a Spatial Transcriptomics dataset generated to study the mouse olfactory bulb.

**Figure 1.**
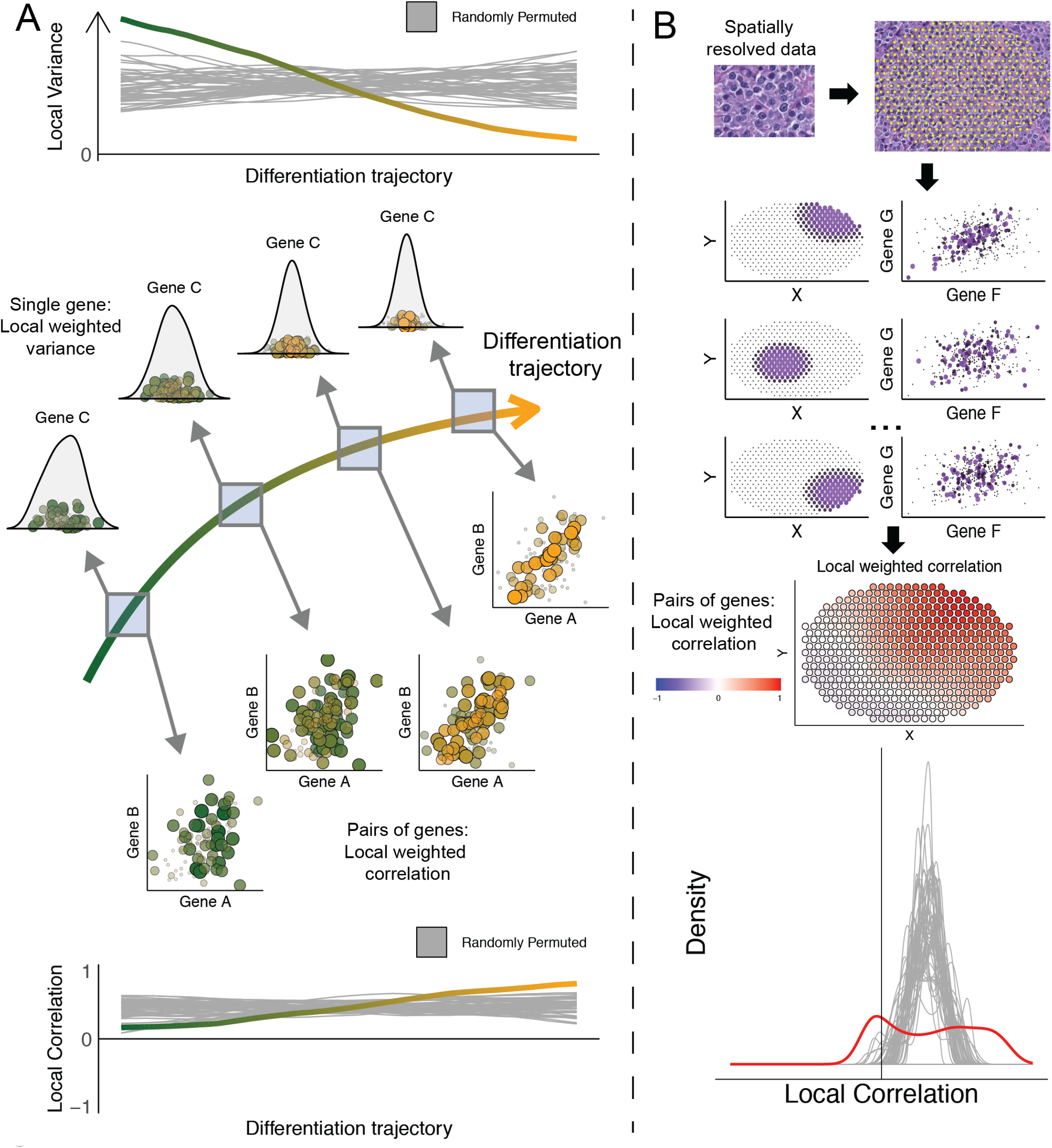
Methods workflow. **A.** Example showing a differentiation trajectory where genes are tested for changes in higher order interactions such as variability and correlation along the trajectory. A set of local higher order statistics are calculated, and significance is compared by repeatedly permuting samples (grey curves). The vector of local estimates of higher order statistics are combined using the sample standard deviation to assess how variable it is across time. **B.** Example showing that in a spatial context, scHOT calculates a field of local estimates of correlation across space, and compares the variability associated with these with permuted sample points across space.

## RESULTS

### scHOT identifies multiple higher order associations during liver development

We first analyzed four single-cell RNA-sequencing datasets designed to study the early development of the mouse liver^27–30^ (Methods, of which three contained hepatic cells). The integrated data encompassed 7 days of development (from embryonic (E) day 10.5 to E17.5), which covers the period where progenitor hepatoblasts transition towards more mature hepatocytes and cholangiocytes (Figure 2A). As expected, when using Monocle^17^ to order the cells in pseudotime, we observed a clear bifurcation where hepatoblasts differentiated into either cholangiocytes or hepatocytes (Figure 2B). This was supported by a higher proportion of differentiated cells in the later embryonic time points as well as cell-type specific expression of known marker genes (Figure 2C).

**Figure 2.**
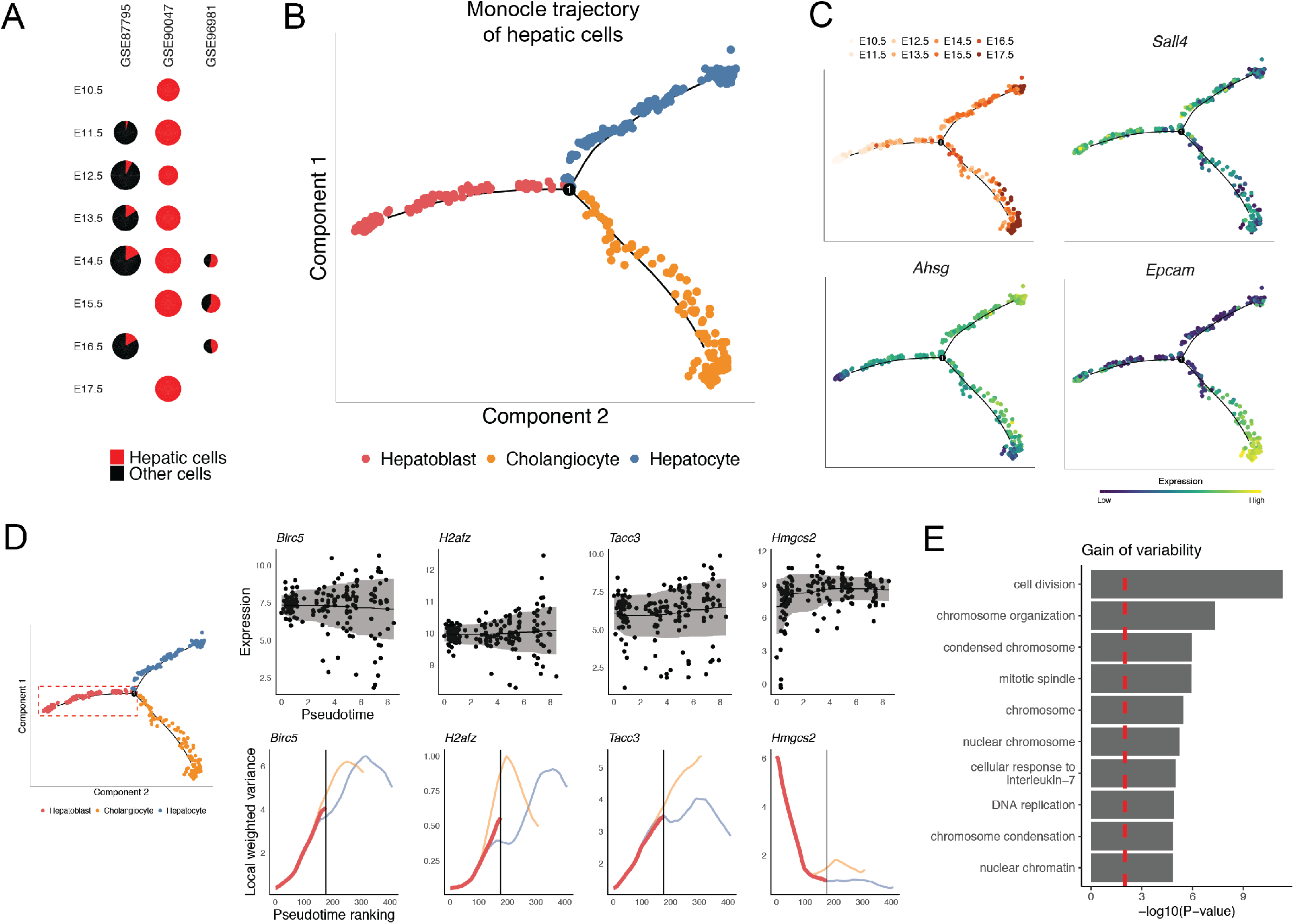
**A.** Charts showing relative proportion of hepatic and non-hepatic cells in the Developmental Liver Data, across original dataset and embryonic stage. **B.** Monocle trajectory of hepatic cells showing a bifurcating trajectory of hepatoblasts into either hepatocytes or cholangiocytes. **C**. Panel of embryonic stage for each cell along the differentiation trajectory, and gene expression of markers of each cell type. **D.** A selection of genes significantly associated with a change in variability along the first branch of the differentiation trajectory, scatterplots show the expression of genes against the pseudotime estimates, and line plots show the local estimate of variance for the first branch (thick lines) and further for the two branches (thin lines). Examples are shown of genes that increase in variability via ‘fanning’ of gene expression along the trajectory (*Birc5*), by a skewed distribution arising (*H2afz*), and by changes in the modality of the expression (*Tacc3* from unimodal to bimodal). *Hmgcs2* is an example of a gene that decreases in variability. **E.** Functional enrichment analysis of genes that significantly increase in variability along the first branch.

Building on this, we first examined higher order patterns as cells transitioned from naïve hepatoblasts towards the bifurcation point where they commit to one of the two downstream lineages (Figure 2D). In total 68 genes showed a change in variability (false discovery rate (FDR) adjusted P-value < 0.1, all shown in Supplementary Figures S2C and S2D) along the trajectory independent of changes in gene expression. Of these, 58 (85%) displayed significantly increased variability along the branch (Figure 2D). These genes were enriched for involvement in processes associated with cell division, chromosomal organization and DNA replication (Figure 2E; Methods), consistent with the notion that increased plasticity can precede cell fate commitment^6,7^.

We next focused on the full trajectory from naïve hepatoblast through to hepatocytes. Specifically, we investigated whether scHOT could identify changes in correlation, independent of differential expression, thus providing insight into the potentially complementary set of gene regulatory modules that are activated during the process of commitment from hepatoblast to the hepatocyte lineage. Correlation patterns identified as cells transition from naive hepatoblast to cholangiocytes can be found in Supplementary Figures S3B and S3C.

When focusing on the hepatoblast to hepatocyte lineage, we identified numerous changes in correlation between pairs of genes that did not change their individual mean expression (Methods). An example of such a gene-pair is *Cdt1* and *Top2a* (Figure 3A, FDR adjusted P-value < 0.03), which are protein-protein interacting partners^31^ that have been implicated in regulation of the cell cycle in human and mouse stem cells^32^. This pair of genes changes from being strongly negatively correlated in the progenitor population to displaying no correlation in the more differentiated hepatocytes. Interestingly, when considering each gene separately, neither *Cdt1* nor *Top2a* are significantly differentially expressed along the trajectory, or significantly differentially variable (FDR adjusted P-value = 0.70 and 0.12 respectively), indicating that the association between these two genes would not be identified without using scHOT. *Top2a* encodes a DNA topoisomerase, which controls and alters the topologic states of DNA during transcription^33^, while *Cdt1* is a chromatin licensing and DNA replication factor that is required for DNA replication and mitosis^34^. Our observation that these genes move from being negatively correlated to displaying no correlation suggests a trade-off between chromatin remodeling and transcription at the earlier stages of differentiation, potentially facilitating both proliferation and the global changes in gene regulatory architecture that arise when cells commit towards the hepatoblast lineage.

**Figure 3.**
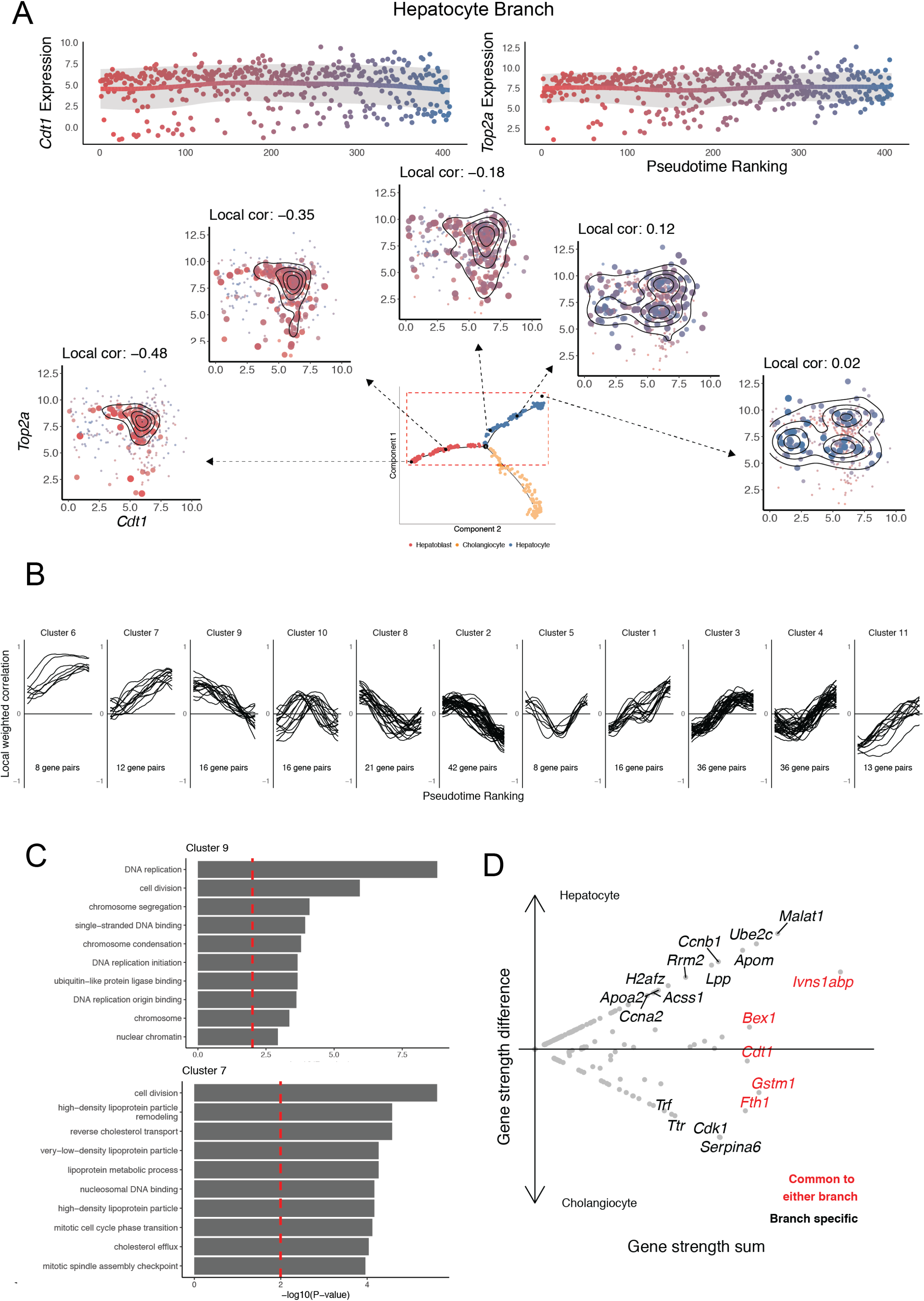
**A.** Sequence of scatterplots showing expression of *Cdt1* and *Top2a* at equally spaced points along the entire trajectory from hepatoblast to hepatocyte. Points are colored by their position along the trajectory, and point size corresponds to the weight given to that region of the trajectory. Neither gene is significantly differentially expressed or differentially variable along the trajectory, but the gene-pair is significantly differentially correlated. **B.** Clustering of local weighted correlation of all significant gene-pairs for the hepatocyte branch, showing groups of gene-pairs that appear to gain or lose correlation across the trajectory. **C.** Functional enrichment analysis of genes belonging to the set of gene-pairs among Clusters 9 and 7 respectively for the hepatocyte branch. **D.** Comparison of hepatocyte and cholangiocyte branches using network strength across all significant gene-pairs for either branch. Black labelled genes are significantly branch specific, while red labelled genes are significantly common across both branches (FDR-adjusted P-value < 0.05).

Across all gene-pairs tested, 224 displayed different patterns of correlation (FDR adjusted P-value < 0.2), encompassing 136 unique genes. Gene-pairs that were differentially correlated were not found to be associated with genes that were also differentially variable along the trajectory (Fisher’s Exact Test P-value > 0.4) for either hepatocyte or cholangiocyte branch, suggesting an independent relationship between changes in correlation of gene-pairs and variability of the genes. The majority of local correlation patterns of these gene-pairs exhibited either a ‘gain’ or a ‘loss’ of correlation, across developmental time reflecting the prior understanding of a continuous differentiation towards the end fate (Figure 3B). Functionally annotating these clusters revealed that, in general, clusters associated with a loss of correlation (e.g., Cluster 9) contained genes linked with DNA replication and cell division. By contrast, clusters that ‘gained’ correlation (e.g., Cluster 7) along the trajectory were associated with hepatocyte-linked functions such as lipoprotein particle remodeling, lipoprotein metabolism, as well as mitotic cell cycle and cell division (Figure 3C, all clusters shown in Supplementary Figure S3A). A small number of gene-pairs displayed more unexpected correlation patterns, with a transient peak of co-expression at an intermediate point along the trajectory (e.g., Clusters 5 and 10) near the bifurcation point, suggesting a transient role in cell fate commitment.

Finally, we explored whether the differential patterns of higher order interactions were associated with specific differentiation into the hepatocyte or cholangiocyte lineages, or a common pattern associated with maturation of cells. To do this, we used the network strength metric as a statistic to assign genes as common to either branch, or branch specific (Methods). Permuting over the topology of the gene network allowed assessment of statistical significance, revealing five genes that were associated with both branches, and ten and four genes significantly specific to the hepatocyte and cholangiocyte branches respectively (Figure 3D). In particular, the gene *Cdt1* appears associated to differentiation in general, i.e. from the hepatoblast state to either differentiated state, rather than any of the two terminal states. By contrast, genes like *Apom* and *Apoa2*, encoding apolipoprotein, were more associated with hepatocyte function. More surprisingly, we identified histone gene *H2afz* as more specific to the hepatocyte lineage, indicating a potential association with changes in global chromatin organization as cell’s commit towards a hepatocyte fate.

### scHOT identifies local patterns of correlation in the mouse olfactory bulb

Finally, we considered whether scHOT could also be used to identify cryptic local correlation when gene expression information is available in a spatial context. Specifically, to date, most studies of spatially resolved gene expression data have focused on clustering cells into groups or testing known patterns of correlation – we reasoned that scHOT would provide an unbiased approach for identifying local patterns of correlation that might be missed by such approaches, which rely on changes in mean expression only. To this end we considered the mouse olfactory bulb (MOB), which displays a highly stereotypical structure, with clear patterns of concentric layers corresponding to granule, internal plexiform, mitral, external plexiform, glomerular and olfactory nerve layers moving from the inside out (Figure 4A), along with distinct patterns of gene expression along this space^35^. Recently, spatial transcriptomics^10^ was used to measure gene expression levels in small spatially-distinct regions of the MOB, thereby facilitating the unbiased identification of patterns of gene expression in space.

**Figure 4.**
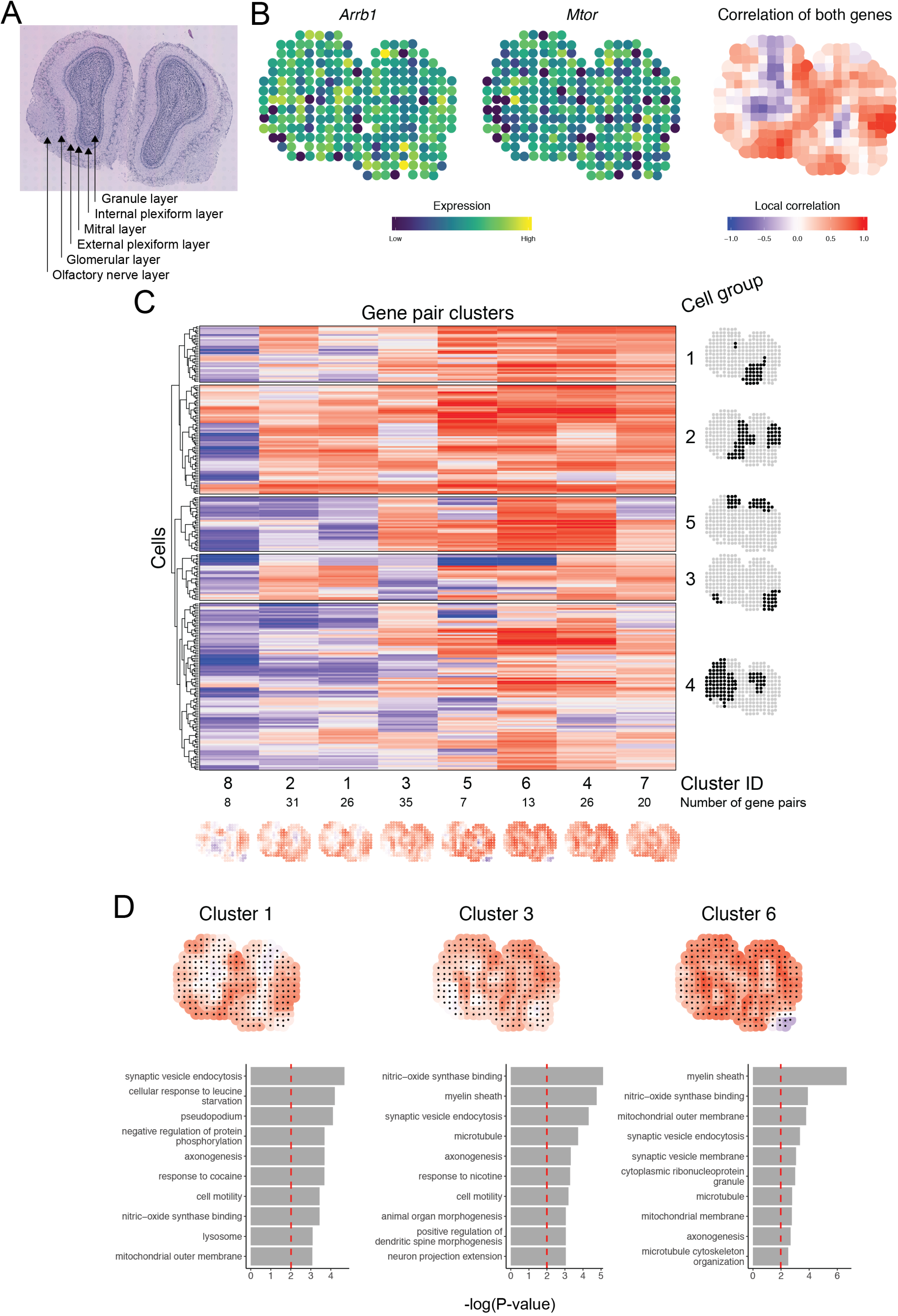
**A.** H&E image of mouse olfactory tissue section with labeling of known anatomical layers. **B.** Spatial expression plots of two genes, *Arrb1* and *Mtor*, which are not significantly differentially expressed across space, and are significantly differentially correlated across space. The third plot shows the local spatial correlation estimated for these two genes, recapitulating the layered pattern of the olfactory bulb. **C**. Heatmap showing all sampled points (rows) and clusters of significantly spatially differentially correlated gene-pairs (columns), with spatial maps of mean local correlation (bottom row) for each group, and highlighted positions (column on right) for the sampled points grouped into clusters. **D.** Spatial maps of local correlation for selected clusters of gene-pairs and barplots showing functional enrichment analysis.

Using scHOT, we identified a set of 167 gene-pairs as significantly differentially correlated across space (FDR adjusted P-value < 0.2), with 42 non-differentially expressed highly variable genes appearing at least once among this set (Methods). Interestingly, we found that numerous pairs of genes displayed diffuse patterns of local correlation that were not apparent when visually comparing their individual expression profiles. For example, *Arrb1* (beta-arrestin-1) is widely expressed in the brain^36^, consistent with its diffuse expression across the MOB in the spatial transcriptomic data. Similarly, *mTOR* (mammalian target of rapamycin), *Uchl1* (Ubiquitin carboxyl-terminal esterase L1), and *Dnm3* (Dynamin-3) are all broadly expressed^37–39^. Nevertheless, we identified all three genes as positively spatially correlated with *Arrb1* (Figure 4B and Supplementary Figure S4C). Consistent with this correlation, *Arrb1* can regulate *mTor* activation^40^ and interact with the ubiquitination pathway to down regulate receptor signaling^41^. Further, both *Arrb1* and *Dnm3* function in endocytosis^42^.

To explore more general patterns, we clustered all significant gene-pairs using their local spatial correlation patterns into 8 distinct groups (Figure 4C). Despite the relatively low-resolution of the data (spatial transcriptomic data is limited to a resolution of ~10 cells (approximately 100 μm)^10^), a variety of local correlation patterns were observed, often associated with distinct biologically meaningful regions of the bulb. Giving confidence in our analysis approach, clustering the cells based on their local correlation pattern largely recapitulated the symmetry of the MOB, with multiple cell groups corresponding to symmetric sets of cells, e.g. Cell groups 3, 4, and 5 (Figure 4C). Additionally, Cluster 1 contained genes that were positively correlated within the Olfactory Nerve Layer and Clusters 3 and 6 were associated with the Mitral and External plexiform layers (Figure 4D). Functional annotation of the genes belonging to these clusters revealed associations with distinct neuronal terms including signaling events such as endocytosis, phosphorylation and ubiquitination (all clusters shown in Supplementary Figure S4D). Interestingly, “myelin sheath” was highly ranked in multiple clusters (Clusters 3-7). In these clusters, the strongest patterns of correlated spatial expression occurred within more internal layers of the bulb, overlapping with the mitral and granule layers. This is consistent with the myelination of the lateral olfactory tract as it exits the bulb^43^. Clusters 1, 2, and to some extent 8, in contrast, possess spatial correlation patterns that encompass more external layers such as the olfactory nerve layer, and genes within these clusters are not highly associated with the term ‘myelin sheath.’ This is consistent with the fact that olfactory sensory neurons entering the bulb in the more external olfactory nerve layer are not myelinated. In sum, we have shown here that exploiting higher-order structure can reveal unexpected and spatially-coherent regions of structured heterogeneity that are independent of mean expression changes, and that approaches that focus only on the latter will fail to fully exploit the wealth of information contained within such data.

## DISCUSSION

In this paper we have demonstrated the utility of higher order testing for single cell data. We examined scHOT in the context of two biological systems with distinct data characteristics – liver development and the mouse olfactory bulb. scHOT is robust due to the choice of underlying higher order metrics such as rank-based Spearman correlation; powerful as it uses a permutation framework retaining the global variability and covariance structure for inference; and extremely flexible as it can be tuned by 1) Varying the local weighting scheme in terms of shape (triangular, block, any other user defined weight) and span, 2) Choice of underlying higher order effect function (weighted Pearson correlation, weighted Spearman correlation, weighted zero-inflated Spearman or Kendall correlation, or any choice of higher order estimate when using the block weighting scheme), and 3) Choice of summarization estimate for the local higher order vector (by default the standard deviation). In general, this contrasts with other methods that estimate changes in expression across either a pseudotime trajectory or across space, which require a set of candidate hypotheses to test explicitly. In the spatial context, scHOT differs substantially to other methods such as SpatialDE^25^, in that we can test either a single gene (identifying spatially variable) or two genes (identify spatially differentially correlated), and no prior suite of potential spatial structures need be provided to identify genes that are of interest.

From a biological perspective, the concept of characterizing coordinated changes over time could enable better characterization of the molecular processes underpinning cell fate choice. In particular, it will help us to better understand whether increased plasticity, as manifested in increased cell-to-cell variability, is a general feature that precedes cell commitment or whether this is restricted to specific systems such as hepatoblast differentiation. Such heterogeneity could also be a driver of differential cell fate or cell function in a spatial context: specific patterns of local correlation could indicate that a specific region of a tissue or organ is primed towards a specific fate. Intriguingly, our reanalysis of data from the mouse olfactory bulb identified patterns that were independent of changes in mean expression but associated with known spatial structure of the bulb.

In summary, scHOT is a method for inference of changes in higher order interactions, not just changes associated with the mean, and as such offers a new lens to interrogate single cell data and describe patterns of variation and covariation, offering additional and complementary insight to that obtained by examining changes in mean expression. It is enabled by a statistical framework that captures nonlinear changes in correlation structure and that uses sample ranking approaches to avoid having to discretize responses. This is especially important for continuous single cell trajectories and for studying spatial structure within ostensibly homogeneous cell types. By facilitating such analysis, scHOT will enable investigations into how highly localized patterns of variation and co-variation influence cell fate and cell function.

## MATERIAL AND METHODS

### Datasets

The following datasets were used to examine scHOT, and demonstrate its utility in extracting insights from diverse sources of single cell and/or spatially resolved data.

#### Developmental Liver Data

The ‘Developmental Liver Data’ is a full-transcript scRNA-Seq dataset generated using plate-based protocols from four distinct sources^27–30^, across multiple mouse embryonic time points from Embryonic Day (E)10.5 to E17.5. The data were integrated using scMerge^44^, taking advantage of genes that are found to be stably expressed in single cells^45^. These data comprise several cell types including hepatic cells such as hepatoblasts, cholangiocytes, and hepatocytes, among other cell types such as immune cells (Figure 2A). Monocle was used to infer a differentiation trajectory exclusively for the hepatic cells. We applied scHOT to these data, considering the following testing scenarios: changes in variability across the first branch from hepatoblasts to the cell fate decision point; and changes in correlation between pairs of highly variable genes along the entire branch from hepatoblasts to hepatocytes and the full branch from hepatoblasts to cholangiocytes.

To select genes for downstream analysis, we considered genes that were highly variable (HVGs)^46^ (Supplementary Figure S2A) but that had consistent mean expression along the trajectory. To do this, we performed liberal differential gene expression testing along pseudotime by fitting, for each gene, two linear models (slope and intercept, and polynomial of degree two) and identifying a gene as differentially expressed if it was significant (F-test; unadjusted P-value < 0.05) in at least one of the tests when compared to an intercept only model. This differential expression testing was performed for the hepatoblast to hepatocyte trajectory and for the hepatoblast to cholangiocyte trajectory. The resulting sets of genes (i.e., highly variable and non-differentially expressed) were combined to form all pairwise combinations as the scaffold for higher order gene-pair testing.

#### Spatial data from the mouse olfactory bulb (MOB)

The ‘Mouse Olfactory Bulb’ data is a Spatial Transcriptomics dataset, where an array spotted with probes that have barcodes corresponding to defined locations was used to measure spatially-resolved gene expression levels^10^. We consider data where this technology was used to measure expression levels across a section of the mouse olfactory bulb (MOB), where each spatially resolved region contains a measure of gene expression averages across approximately 10 cells^10^. This cross section of the MOB comprises concentric layers visible with H&E staining (Figure 4A), associated with the granule, internal plexiform, mitral, external plexiform, glomerular and olfactory nerve layers moving from the inside out. The resulting expression data is derived from high throughput RNA sequencing using barcodes corresponding to the spatial locations, as well as unique molecular identifier (UMI) barcodes. We identified genes as spatially differentially expressed by performing scHOT using a first-order metric of local weighted mean expression (2,542 differentially expressed genes; unadjusted P-value < 0.05). After identifying the intersection between genes that were highly variable^46^ but not differentially expressed we used scHOT to test all pairwise combinations for this set of genes.

### Choice of local weighting scheme

For the trajectory-based analyses we selected a triangular weight matrix with a span of 0.25. For the spatial-based analysis we selected a two-dimensional triangular weight matrix (i.e. a cone) also with a Euclidean distance span of 0.05 (here corresponding roughly to 9 surrounding sampled points).

### scHOT test statistic and inference

For single gene testing we use a local weighted variance estimate

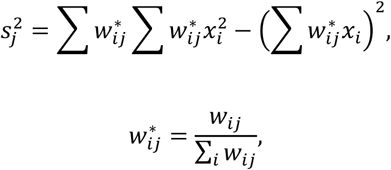

where *w*_*ij*_ is the cell-specific weight for cell *i* and position *j*, and *x*_*i*_ is the gene expression measure for gene *x* and cell *i*. For testing pairs of genes we use a weighted Spearman correlation

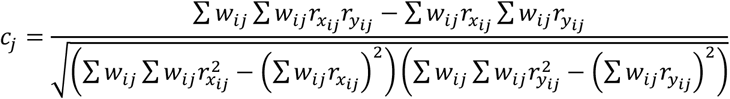

where

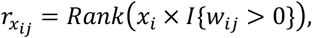

and

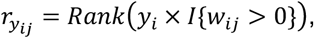

where additionally *y*_*i*_ is the gene expression measure for gene *y* and cell *i*.

The scHOT test statistic is a measure of variability associated with this vector of local variances or correlations. To compute this, we first calculate the sample standard deviation to estimate the variability associated with the set of local variance estimates 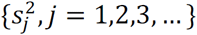 or local correlation estimates {*c*_*j*_, *j* = 1,2,3, …}.

Statistical testing is then performed by randomly permuting cell labels, while keeping the overall gene expression structure constant. Thus, within each permutation round, the global correlation or global variance remains the same, while the vectors of local variability or local correlation vary. In all cases, we used sample or cell permutation and defined significance by controlling for a 0.2 Benjamini-Hochberg^47^ FDR in all differential correlation tests, and at 0.1 FDR for variability tests. For correlation-based tests, we used the fact that the null distribution is based only on the two matched gene expression vectors to interpolate null distributions given the global correlation value and the number of samples (see Computational efficiency section) in order to speed up computation. In the case of discrete groupings, we use a normal approximation to the null distribution to estimate high resolution p-values.

### Downstream analysis

After identifying significant gene-pairs, we took their local correlation profiles and hierarchically clustered them to identify patterns of differential correlation across either pseudotime or space, using maximum distance and Ward’s minimum variance method. For the Liver Developmental Dataset correlation analysis, we smoothed local correlation vectors before hierarchical clustering using loess. For both datasets we extracted discrete clusters from the hierarchical clustering using the R function cutree with number of clusters estimated using dynamicTreeCut^48^.

### Comparing between trajectory branches

We implemented a statistical test for comparing the change in correlation between the two branches, by examining the normalized network strength across the tested networks per branch. We defined network strength for a given gene (node) as the sum of edge weights for significant gene-pairs associated with the gene, divided by the total edge weights across the entire network. The edge weight we selected was the −log(FDR-adjusted P-value) for each gene-pair. For each gene, we calculated the network strength of all genes per branch. We then compared these network strength values between branches using an MA-plot, i.e. comparing the sum of network strengths with the difference of network strengths. To assess significance associated with a single gene – i.e. a gene that tends to have a higher network strength than expected by chance, we repeatedly permuted the gene-pair edge weights across the network and calculated the permuted MA-plot. Individual genes were identified with a significantly nonzero network strength using the Euclidean distance from the origin as the test statistic. To identify genes with a branch-specific network strength, we considered the ratio of significance towards each branch as the test statistic.

### Computational efficiency

We previously observed a relationship between the total number of samples and the null distribution of the DCARS test statistics^24^. Here we uncovered further association between the null distribution of the scHOT test statistics and the global correlation across all samples. This represents an opportunity to ignificantly decrease computational time as one can ‘borrow’ permutations from similarly distributed genes and gene-pairs to estimate the p-value. Our approach is to first calculate global correlations for all gene-pairs to be tested, and then take a uniform sample among the gene-pairs according to the global correlations. For this subset of gene-pairs we permute sample labels and calculate scHOT test statistics. Then for any given gene-pair of interest, we extract the desired number of permutations from this set of permuted scHOT test statistics, according to how similar their global correlation is. These are shown in Supplementary Figure S2E and S4B for the liver and MOB data respectively.

## DATA AND SOFTWARE AVAILABILTY

All data analysis was performed on publicly available data. The liver developmental dataset and description is available from https://sydneybiox.github.io/scMerge/articles/Mouse_Liver_Data/Mouse_Liver_Data.html and the specific R data file downloaded from http://www.maths.usyd.edu.au/u/yingxinl/wwwnb/scMergeData/liver_scMerge.rds

The mouse olfactory bulb data was downloaded from the Spatial Research website https://www.spatialresearch.org/resources-published-datasets/doi-10-1126science-aaf2403/ and count matrix data and H&E stained brightfield image related to MOB replicate 11 was downloaded.

Instructions to access data, software, and scripts to perform the analysis presented here is available at https://github.com/MarioniLab/scHOT2019.

## ACKNOWLEDGEMENT

The authors thank all their colleagues, particularly at Cancer Research UK Cambridge Institute and The University of Sydney School of Mathematics and Statistics for their support and intellectual engagement. In particular, the authors gratefully acknowledge Pengyi Yang and Michael Morgan for their helpful discussion.

## AUTHOR CONTRIBUTIONS

S.G. conceived the study with input from E.P., J.Y.H.Y. and J.C.M.. S.G. developed the method and performed data analysis with input from Y.L. S.G., D.M.L and X.S. interpreted the results with input from Z.G.H. and J.C.M.. S.G., J.C.M. and J.Y.H.Y. wrote the manuscript. All authors read and approved the final version of the manuscript.

## SUPPLEMENTARY FIGURE LEGENDS

**Supplementary Figure S1.**
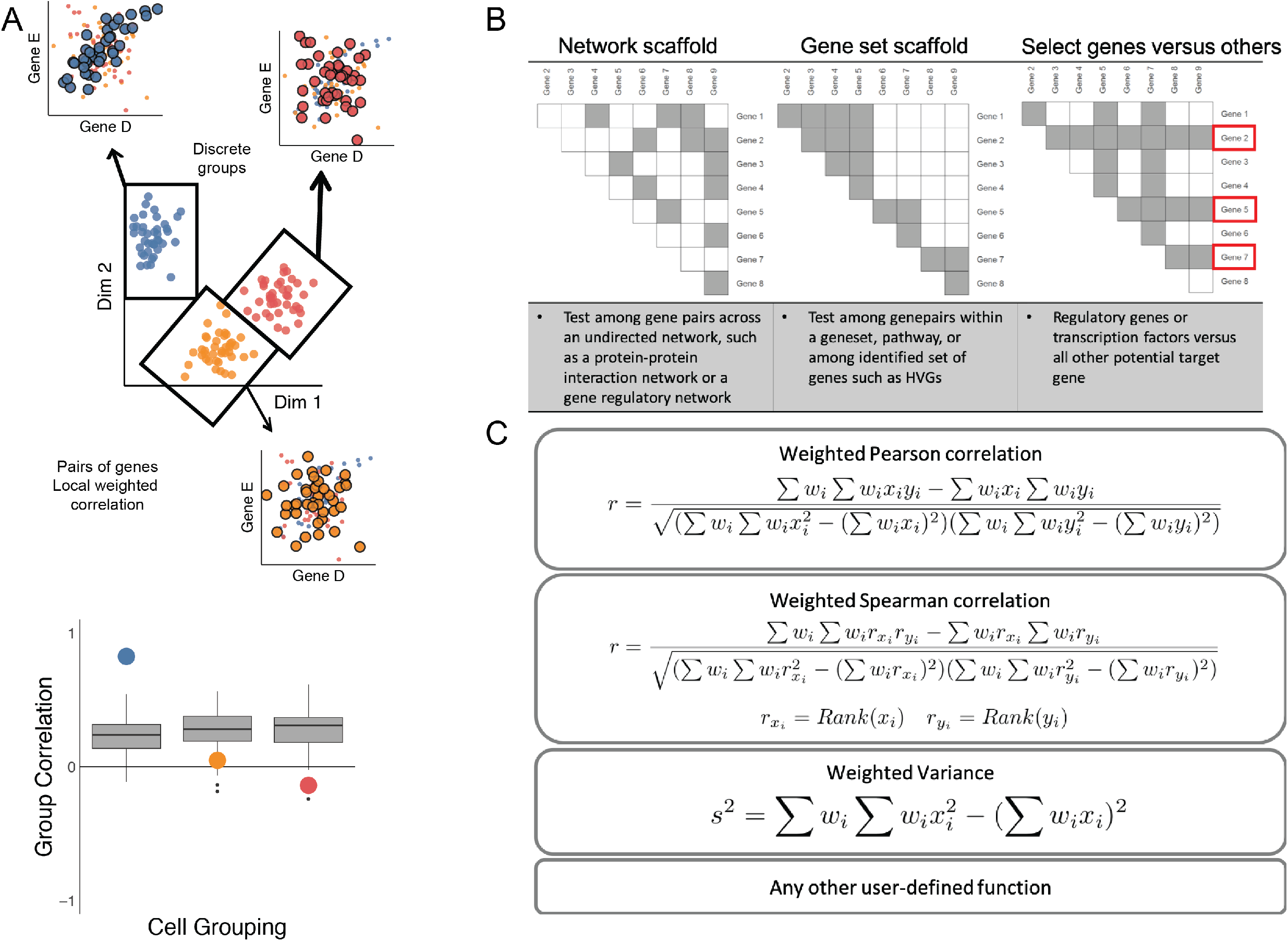
**A.** Example showing testing for correlation differences in three distinct groups. A set of local higher order statistics are calculated, and significance is compared by repeatedly permuting samples (grey boxplots). The set of local estimates of higher order statistics are combined using the sample standard deviation to assess how variable they are between groups. **B.** Possible schemes for the testing scaffold using gene networks, including: i) a gene-gene network; ii) a gene set scaffold where all pairwise combinations within a gene set are included; and iii) selected genes of interest versus all others. **C.** Examples of weighted higher order functions including weighted Pearson correlation, weighted Spearman correlation, weighted variance. Note that any user defined function can be used.

**Supplementary Figure S2.**
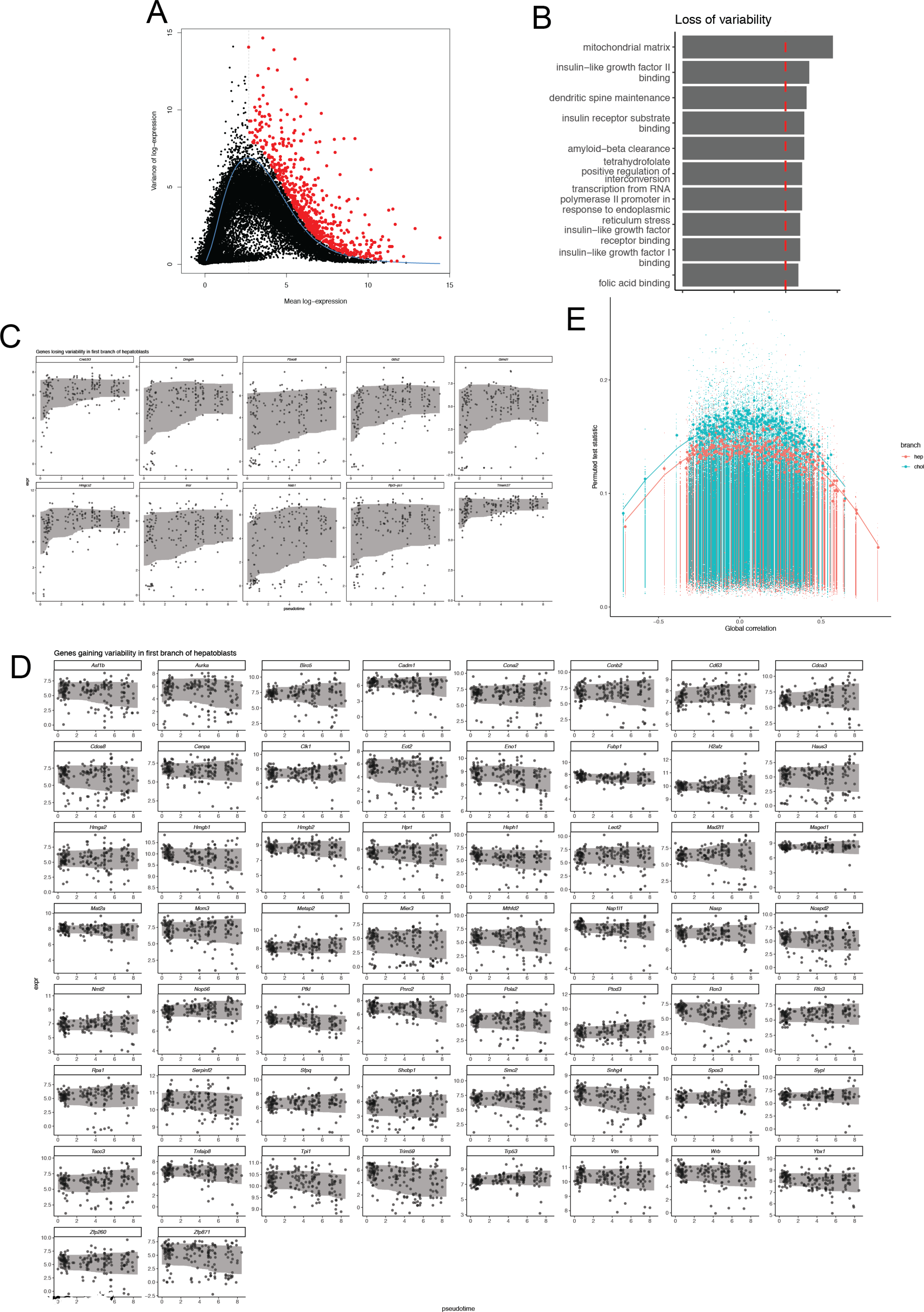
**A.** HVG selection for Developmental Liver Data. **B.** Barplot for Gene Ontology testing for genes showing loss of variability along hepatoblast branch. **C.** Scatter and ribbon plot of genes showing loss of variability along hepatoblast branch. **D.** Scatter and ribbon plot of genes showing gain of variability along hepatoblast branch. **E.** Global correlation and null scHOT correlation test statistics for both hepatocyte and cholangiocyte branches.

**Supplementary Figure S3.**
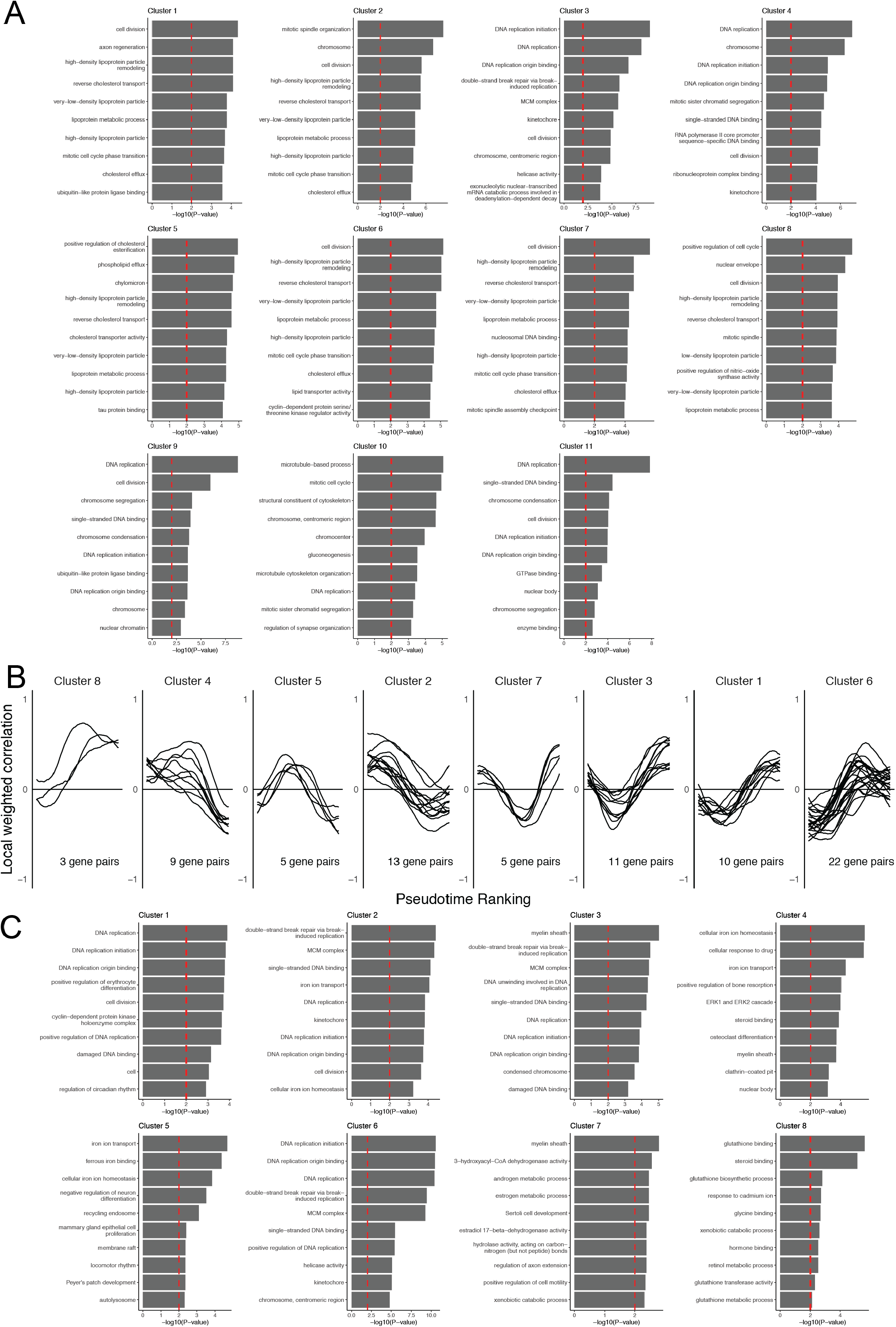
**A.** Gene ontology barplots for all scHOT hepatocyte clusters. **B.** Line plots of clustered significant scHOT gene-pairs for full cholangiocyte branch. **C.** Gene ontology barplots for all scHOT cholangiocyte clusters.

**Supplementary Figure S4.**
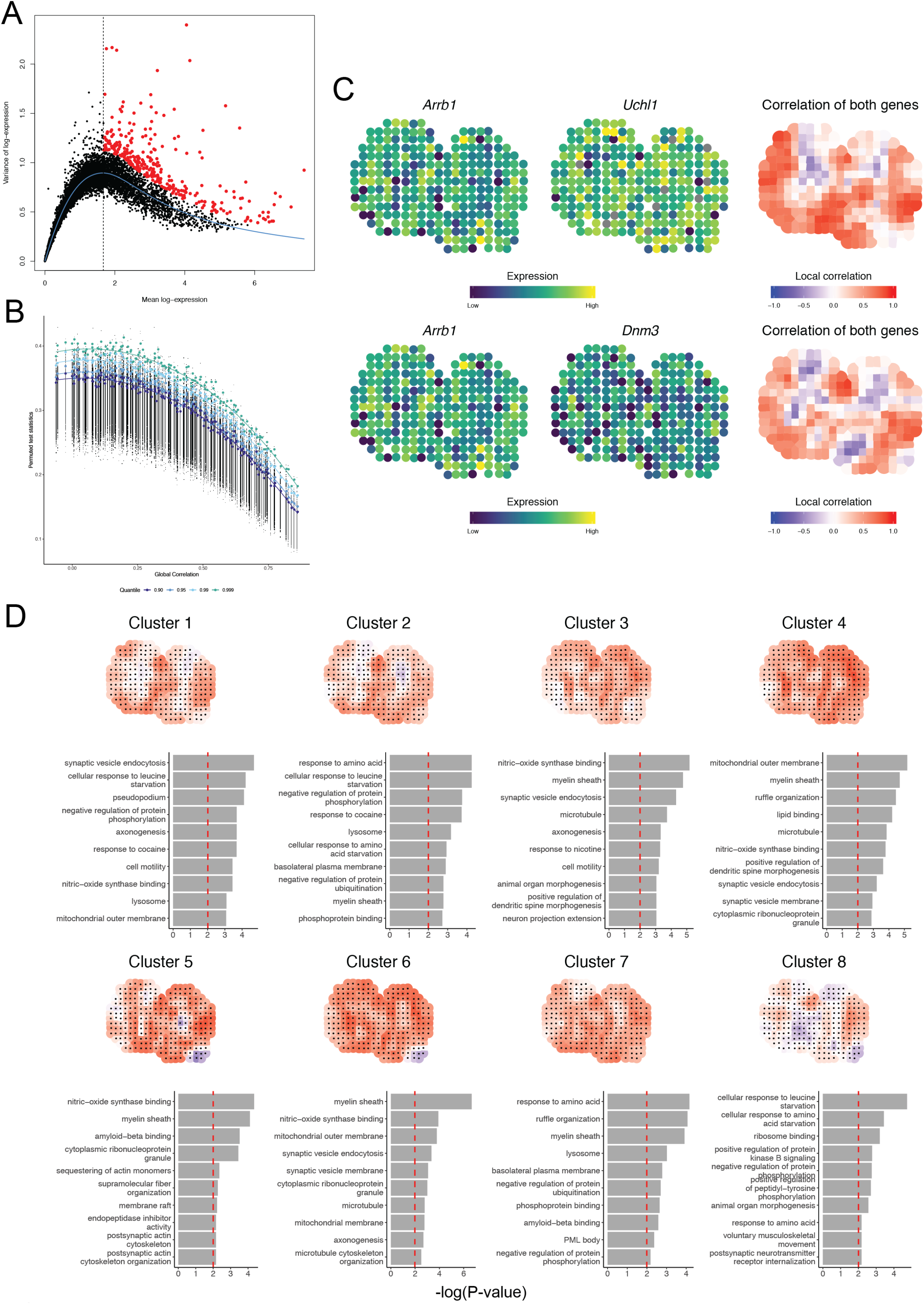
**A.** HVG selection for Spatial Transcriptomics analysis. **B.** Global correlation and null scHOT correlation test statistics for spatial MOB data. **C.** Spatial expression plots of two gene-pairs *Arrb1* and *Uchl1* as well as *Arrb1* and *Dnm3* which are not significantly differentially expressed across space, but are significantly differentially correlated across space. The third plot shows the local spatial correlation estimated for these two genes, recapitulating the layered pattern of the olfactory bulb. **D.** Spatial maps of mean local correlation and gene ontology functional enrichment barplots for all MOB scHOT clusters.

## SUPPLEMENTARY TEXT

### scHOT extensions and considerations

scHOT is a flexible framework within which multiple aspects can be modified to facilitate bespoke analysis. For example, higher order patterns can be studied along trajectories, across space, or among discrete groups (Supplementary Figure S1A). Moreover, distinct sets of genes or gene-pairs can be interrogated depending on the biological question of interest (Supplementary Figure S1B). Of particular interest, the local weighting scheme and concordance function can also be adapted, depending upon the biological context (some examples are given in Supplementary Figure S1C). For example, if one were interested in identifying changes in higher order interactions along a circular trajectory, e.g. the cell cycle, one could define a local weighting scheme that was also circular – by ensuring that the two ends match given any starting point. Another example is for discrete groups that are either completely distinct, or ordered in some way – e.g. over discrete time points along a time course experiment, one may define the weight matrix to incorporate the discrete grouping, while also accounting for the flanking groups. More generally, one may wish to place a higher local weight over a particular local region and a smaller weight over a different region. We note, however, that these changes to the weighting scheme may affect the generalizability of the null distributions across multiple genes or gene-pairs, so the user should take care in ensuring that the null distributions appear similar when employing computational speed-up steps.

Any concordance metric can be ‘plugged in’ if using a block type of weight matrix. That is, one is able to use the fast implementation of distance correlation between two distributions^49^, mutual information, partial correlation, or any concordance metric suited especially to other data types such as ordinal or binarized single cell data^50^, without needing to explicitly derive weighted formulations of these concordance metrics. Additionally, any concordance metric that doesn’t necessarily have a ‘weighted’ formulation and/or implementation can be utilized using block weighting scheme. This makes scHOT versatile, by enabling user-defined metrics. For summarizing the vector of local higher order statistics, users may wish to substitute the sample standard deviation with any other choice of variability or change estimate – e.g. if the goal is to examine how monotonic a change in higher order interaction is, a measure of monotonicity such as mutual information or Spearman correlation with the weighting scheme index could be used.

